# (p)ppGpp and DksA play crucial role in reducing the efficacy of β-lactam antibiotics by modulating bacterial membrane permeability

**DOI:** 10.1101/2024.05.08.593222

**Authors:** Meenal Chawla, Jyoti Verma, Shashi Kumari, Tushar Matta, Tarosi Senapati, Prabhakar Babele, Yashwant Kumar, Rupak K Bhadra, Bhabatosh Das

## Abstract

The key signaling molecules in the bacterial stress sensing pathway, the alarmone (p)ppGpp and transcription factor DksA, help in survival during nutritional deprivation and exposure to xenobiotics by modulating cellular metabolic pathways. In *Vibrio cholerae*, (p)ppGpp metabolism is solely linked with the functions of three proteins: RelA, SpoT, and RelV. At threshold or elevated concentrations of (p)ppGpp, the level of cellular metabolites and proteins in the presence and absence of DksA in *V. cholerae* and other bacteria has not yet been comprehensively studied. We engineered the genome of *V. cholerae* to develop DksA null mutants in the presence and absence of (p)ppGpp biosynthetic enzymes. We observed a higher sensitivity of the (p)ppGpp^0^Δ*dksA V. cholerae* mutant to different ꞵ-lactam antibiotics compared to the wild-type (WT) strain. Our whole-cell metabolomic and proteome analysis revealed that the cell membrane and peptidoglycan biosynthesis pathways are significantly altered in the (p)ppGpp^0^, Δ*dksA*, and (p)ppGpp^0^Δ*dksA V. cholerae* strains. Further, the mutant strains displayed enhanced inner and outer membrane permeability in comparison to the WT strains. These results directly correlate with the tolerance and survival of *V. cholerae* to ꞵ-lactam antibiotics. These findings may help in the development of adjuvants for ꞵ-lactam antibiotics by inhibiting the functions of stringent response modulators.

**Importance:** The (p)ppGpp biosynthetic pathway is widely conserved in bacteria. Intracellular levels of (p)ppGpp and the transcription factor DksA play crucial roles in bacterial multiplication and viability in the presence of antibiotics and/or other xenobiotics. The present findings have shown that (p)ppGpp and DksA significantly reduces the efficacy of ꞵ-lactam and other antibiotics by modulating the availability of peptidoglycan and cell membrane-associated metabolites by reducing membrane permeability. Nevertheless, the whole-cell proteome analysis of (p)ppGpp^0^, Δ*dksA*, and (p)ppGpp^0^Δ*dksA* strains identified the biosynthetic pathways and associated enzymes that are directly modulated by the stringent response effector molecules. Thus, the (p)ppGpp metabolic pathways and DksA could be a potential target for increasing the efficacy of antibiotics and developing antibiotic adjuvants.

## INTRODUCTION

The stringent response (SR) is a predominant bacterial response to stress consisting of inadequate nutrients, which permits bacteria to swiftly adjust to their immediate environment. SR is characterized by major cellular reprogramming which includes the downregulation of stable RNA (rRNA and tRNA) synthesis, upregulation of amino acid biosynthesis, and other central metabolisms crucial for survival (1–4). SR is mediated by two small intracellular signaling molecules, guanosine 3’-diphosphate 5’-triphosphate (pppGpp) and guanosine 3’, 5’- (bis) diphosphate (ppGpp), collectively known as (p)ppGpp (5, 6). The (p)ppGpp has an authoritative regulator of nearly every facet of bacterial physiology, such as growth rate regulation, phase shift, toxin generation, biofilm development, motility, a plethora of additional virulence connections, and antimicrobial resistance (1, 2, 7). The non-DNA binding transcriptional factor DksA is also a SR master regulator (8). DksA was discovered as a multicopy suppressor of a *dnaK* mutant (9), and was later postulated to work at the translational stage, where it was discovered to adhere to RNA polymerase (RNAP) (10, 11).

*Vibrio cholerae,* a well-studied human pathogen and the etiological agent of acute diarrheal disease cholera, is equipped with distinct (p)ppGpp metabolic pathways compared to other Gram-negative bacteria, including the model organism *Escherichia coli* (12). In *V. cholerae*, the regulation of intracellular (p)ppGpp metabolism involves two multidomain proteins, RelA and SpoT, along with a small alarmone synthetase known as RelV (12–14). The (p)ppGpp alone or in combination with DksA interacts with RNAP to regulate transcriptional responses of more than 750 genes in *E. coli* (15, 16). The role of (p)ppGpp and DksA in modulating the antimicrobial susceptibility has been studied in different pathogens (17–22), however, the mechanisms remain unclear.

In this study, we have done extensive genetic engineering on *V. cholerae*, aiming to investigate the impact of the (p)ppGpp and DksA deficits on modulating the antimicrobial susceptibility. To elucidate the connection between these factors, the strains with (p)ppGpp^0^, Δ*dksA*, and (p)ppGpp^0^Δ*dksA* genetic backgrounds were used for antimicrobial susceptibility testing against different antibacterial agents commonly used to manage different infections. To gain insights into the underlying mechanisms, non-targeted metabolomics and proteomics approaches are employed to decode the metabolites and proteins that are directly connected with the stringent response. This integrated analysis aimed to unravel the changes associated with the observed differences in antibiotic susceptibility, shedding light on the role of (p)ppGpp and DksA in the adaptive responses of *V. cholerae* to antimicrobial exposure.

## RESULTS

### (p)ppGpp and DksA reduce the efficacy of antibiotics

Antibiotic susceptibility tests by the disc diffusion method were conducted to measure the zone of clearance with different antibiotics in all the genetically modified strains (Table 1). A total of 14 antibiotics belonging to different classes (Table S1) were used. The level of (p)ppGpp and absence of *dksA* functions are shown to be inversely correlated with the zone of clearance for several antibiotics (Fig. 1). For the majority of the antibiotics, the JV9 strain, a (p)ppGpp^0^Δ*dksA* mutant, was found to be the most sensitive. Also, the strain with higher level of (p)ppGpp, BS1.1, JV8 ((p)ppGpp high:Δ*dksA)* were shown to have considerably lower zone diameters for the β-lactam antibiotics ampicillin and penicillin as compared to BRV1((p)ppGpp^0^) and JV9. The absence of *dksA* gene has especially shown increased sensitivity to erythromycin, a macrolide antibiotic and gentamicin, an aminoglycoside antibiotic (Fig. 1). There is an intrinsic resistance to colistin and polymyxin B in all the strains, and resistance to kanamycin, streptomycin, spectinomycin, chloramphenicol, and zeocin in specific strains was due to the presence of resistance genes inserted during the genome engineering for gene knockout. These antibiotics were not considered for differential sensitivity test.

**Figure 1:**
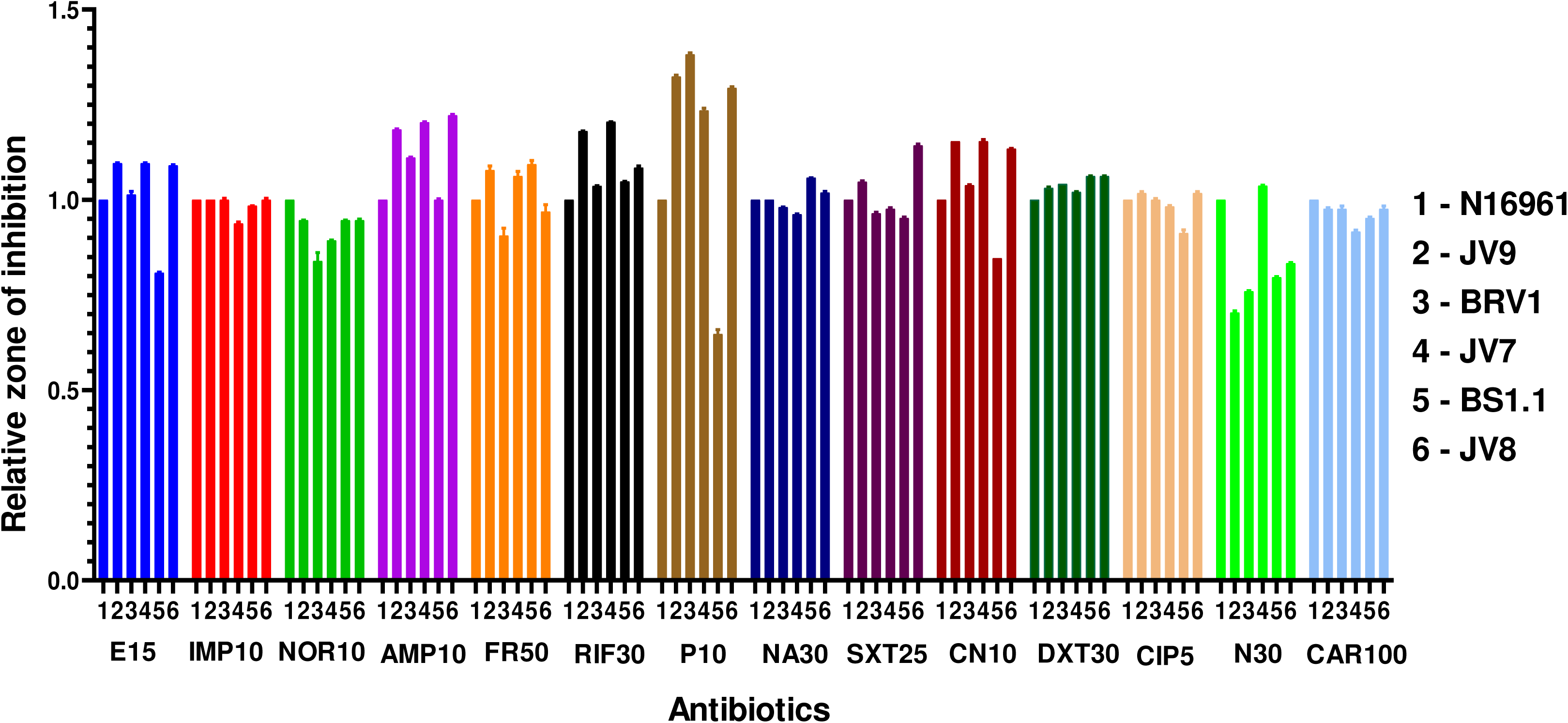
**A)** The graph represents the relative zone of Inhibition of *V. cholerae* WT, JV9, BRV1, JV7, BS1.1 and JV8 strains with different antibiotics.

**Table 1:** Relevant genotype and phenotype of wild-type and genetically modified *Vibrio cholerae* strains used in the study.

| Strain | Genotype and Phenotype | References |
| --- | --- | --- |
| N16961 | Wild type, O1 El Tor, Stp<sup>r</sup> | <u>(27)</u> |
| NR13 | N16961ΔrelA, Stp<sup>r</sup>, Kan<sup>r</sup> | <u>(25)</u> |
| BS1.1 | N16961ΔrelA, ΔspoT, Stp<sup>r</sup>, Kan<sup>r</sup>, Cam<sup>r</sup> | <u>(26)</u> |
| NRV1 | N16961ΔrelV, Stp<sup>r</sup>, Spec<sup>r</sup> | <u>(26)</u> |
| RRV1 | N16961ΔrelVΔrelA, Stp<sup>r</sup>, Spec<sup>r</sup>, Kan<sup>r</sup> | <u>(26)</u> |
| BRV1 | N16961ΔrelVΔrelAΔspoT, Stp<sup>r</sup>, Spec<sup>r</sup>, Kan<sup>r</sup>, Cam<sup>r</sup> | <u>(26)</u> |
| JV7 | N16961ΔdksA, Stp<sup>r</sup>, Zeo<sup>r</sup> | This study |
| JV8 | BS1.1ΔdksA, Stp<sup>r</sup>, Kan<sup>r</sup>, Cam<sup>r</sup>, Zeo<sup>r</sup> | This study |
| JV9 | BRV1ΔdksA, Stp<sup>r</sup>, Cam<sup>r</sup>, Spec<sup>r</sup>, Kan<sup>r</sup>, Zeo<sup>r</sup> | This study |
| MC1 | NR13ΔdksA, Stp<sup>r</sup>, Kan<sup>r</sup>, Zeo<sup>r</sup> | This study |
| MC3 | RRV1ΔdksA, Stp<sup>r</sup>, Spec<sup>r</sup>, Kan<sup>r</sup>, Zeo<sup>r</sup> | This study |
| MC4 | NRV1ΔdksA, Stp<sup>r</sup>, Spec<sup>r</sup>, Zeo<sup>r</sup> | This study |

### (p)ppGpp and DksA augment survival and multiplication of *V. cholerae* in the presence of a sub-lethal dose of the β-lactam

The increased sensitivity of (p)ppGpp and DksA mutant strains to β-lactam antibiotics prompted us to investigate the roles of these factors in the survival of bacteria when exposed to antibiotics in the growth medium. To comprehend their significance, we selected four antibiotics that primarily act on bacterial cell wall biosynthesis. Out of four antibiotics, two antibiotics, ampicillin and vancomycin, exhibited decrease the survival rate of JV9 and JV7 (Δ*dksA)* strains while there were increased rates of survival in the strain with a higher level of (p)ppGpp, BS1.1 and JV8 as compared to WT strain N16961 (Fig 2A-2D). Ampicillin was also shown to be lethal for the MC4 (WT:Δ*relV*Δ*dksA*) strain. None of the bacterial strains had antibiotic resistance genes against these selected antibiotics. Thus, results suggest that the level of (p)ppGpp and presence and absence of DksA are directly correlated with the survival ability of bacterial cells. Different (p)ppGpp synthetases/hydrolases recognized the presence of various antibiotics and translated the signal to (p)ppGpp metabolic pathway to accumulate the alarmone and help *V. cholerae* to sustain in the presence of antimicrobial agents. Similarly, DksA, either independently or in conjunction with (p)ppGpp can modulate transcriptional regulation of various genes involved in metabolic functions directly or indirectly linked to antibiotic resistance.

**Figure 2:**
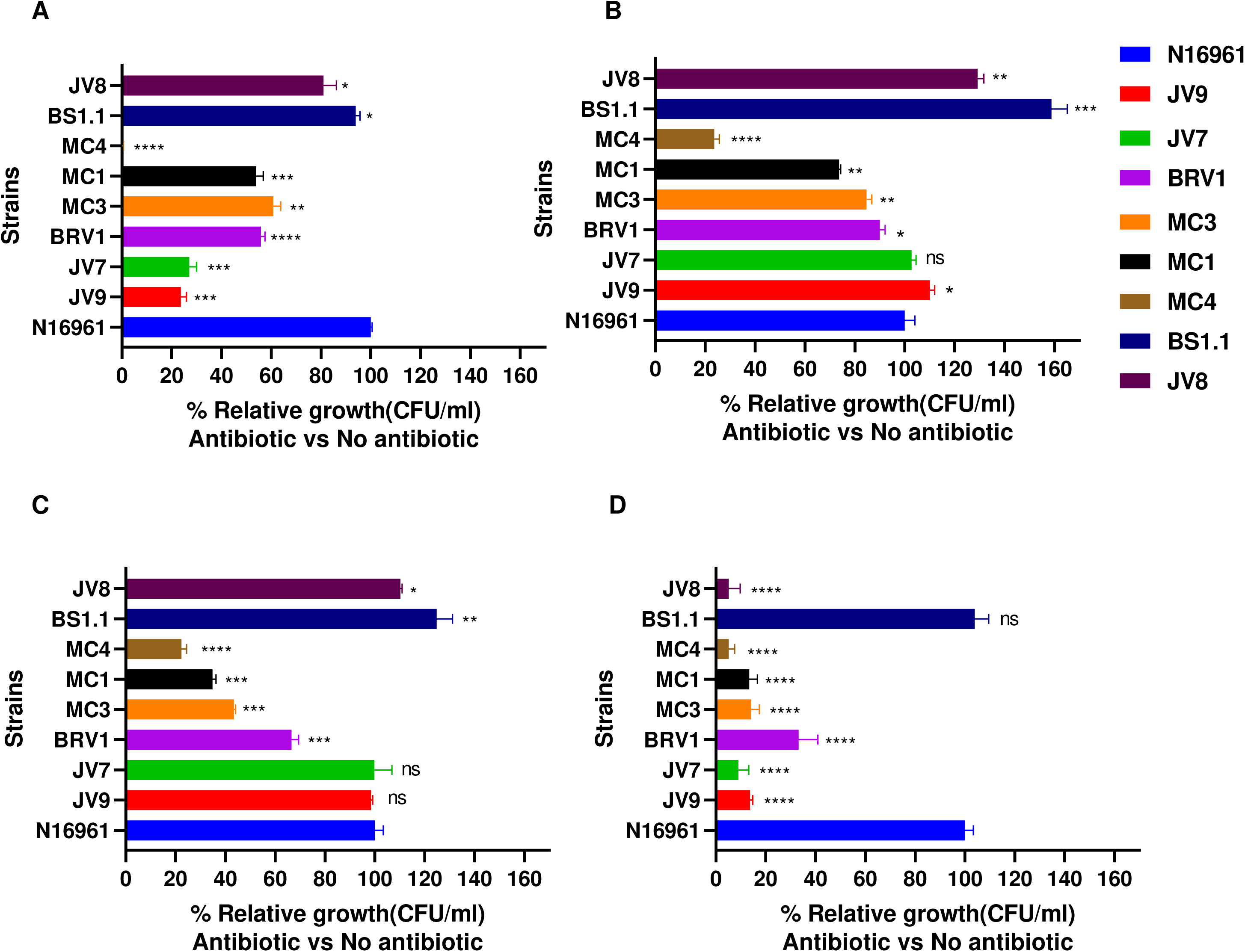
The graph showing the ability of survival of different mutant strains of *V. cholerae* in the presence of **B)** Ampicillin, **C)** Imipenem, **D)** Aztreonam, and **E)** Vancomycin antibiotics. (p-values <0.05 (*), <0.01 (**), p-values <0.001 (***) and p-values < 0.0001 (****)).

### Differential expression of peptidoglycan and cell membrane associated metabolites in (p)ppGpp^0^ and DksA mutants

To unravel the mechanisms contributing to the heightened susceptibility of mutant strains to β-lactam antibiotics, an untargeted metabolomic analysis was conducted. Liquid chromatography coupled mass spectrometry-based profiling was used to study the diversity and abundance of metabolites of genetically engineered *V. cholerae* strains with or without (p)ppGpp synthetase/hydrolase encoding genes and their combination with presence and absence of DksA functions. This comparison is the first metabolomic study to determine the chemical fingerprint of *V. cholerae* (p)ppGpp^0^ and DksA variant strains, as well as the important biochemical pathways involved in the formation of essential metabolites in WT and (p)ppGpp and DksA mutant *s*trains. A total of around 291 metabolites were identified. The combined and individual effects of (p)ppGpp levels and the DksA mutation have been studied, and each mutant strain showed a substantial difference from the WT strain. The PCA plot illustrates the differences between the WT, BRV1, JV7, and JV9 strains (Fig 3A). The significance versus fold-change of all the identified metabolites for the JV9 strain is shown in the volcano plot (Fig 3C). Chemical enrichment analysis was performed in order to provide the chemical classes significantly altered in BRV1, JV7 and JV9 wrt to WT strain individually. The Chemical enrichment analysis of BRV1, JV7 and JV9 strains further corroborated the differential results in the mutant strains (Fig 3D, 3E, and 3F, respectively). Chemical classes with high enrichment ratio for BRV1 include sterol lipids, alkaloids and fatty acyls among others and chemical classes with high enrichment ratio for JV7 strain include carbohydrates, nucleic acids, and glycerophospholipids. Whereas for JV9 strain nucleic acids, carbohydrates, polyketides and lipids chemical classes were having high enrichment ratio.

**Figure 3:**
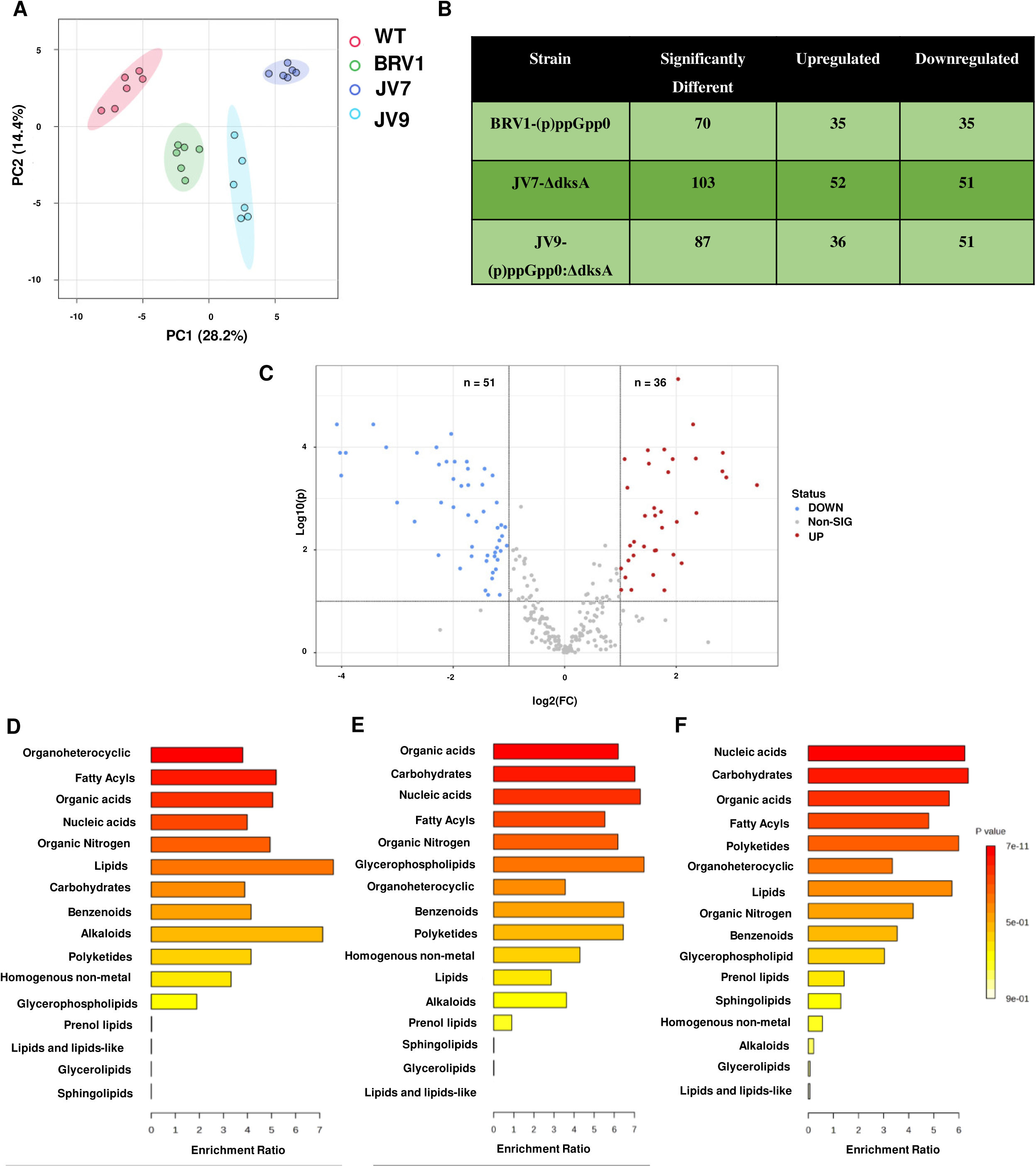
Chemometric analysis of metabolites among (p)ppGpp^0^ (BRV1), Δ*dksA* (JV7), (p)ppGpp^0^:Δ*dksA* (JV9) and WT controls (N16961). **A)** 2D Principal Component Analysis (PCA) score plot of four groups. **B)** Table illustrating metabolites significantly upregulated and downregulated in BRV1, JV7, and JV9 strains, with a fold-change threshold of ±2. **C)** Volcano plot of the 291 identified metabolites of JV9 by LC–MS. The volcano plot shows the fold-change (x-axis) versus the significance (y-axis) of the 291 metabolites. The vertical and horizontal dotted lines show the cut-off of fold-change = ± 2, and of p-value = 0.05, respectively. Metaboanalyst based chemical-enrichment analysis of the identified metabolites in **D)** BRV1, **E)** JV7 and **F)** JV9 strains.

We identified differential patterns in metabolites associated with amino acid metabolism, nucleotide metabolism, and the tricarboxylic acid (TCA) cycle in (p)ppGpp and DksA mutant strains. In addition to elucidating their roles in these key cellular processes, our study unveiled regulatory effects by (p)ppGpp and DksA on metabolites specifically related to cell wall and membrane components. The nucleotide sugar uridine diphosphate N-acetylgalactosamine, also known as UDP-GalNAc, is the precursor for the formation of lipid A of lipopolysaccharide (LPS), which constitutes the majority of Gram-negative bacteria outer membrane (23). The levels of UDP-GalNAc in BRV1 strain were observed to be up-regulated, in contrast no change in UDP-GalNAc levels was observed in BS1.1 strain. The other crucial component of LPS, 2-Keto-3-deoxy octanoic acid (KDO) present in the outermost leaflet of the Gram-negative bacterial outer membrane, was observed to be higher in the BRV1 and JV7 strain compared to WT strain (Fig. 4C). Lipid IVA receives the two Kdo sugar from the core oligosaccharide. The enzyme WaaA, formerly known as KdtA, mediates this step by successively adding Kdo groups to the lipid IVA from activated Kdo (CMP-Kdo)(24–27).

**Figure 4:**
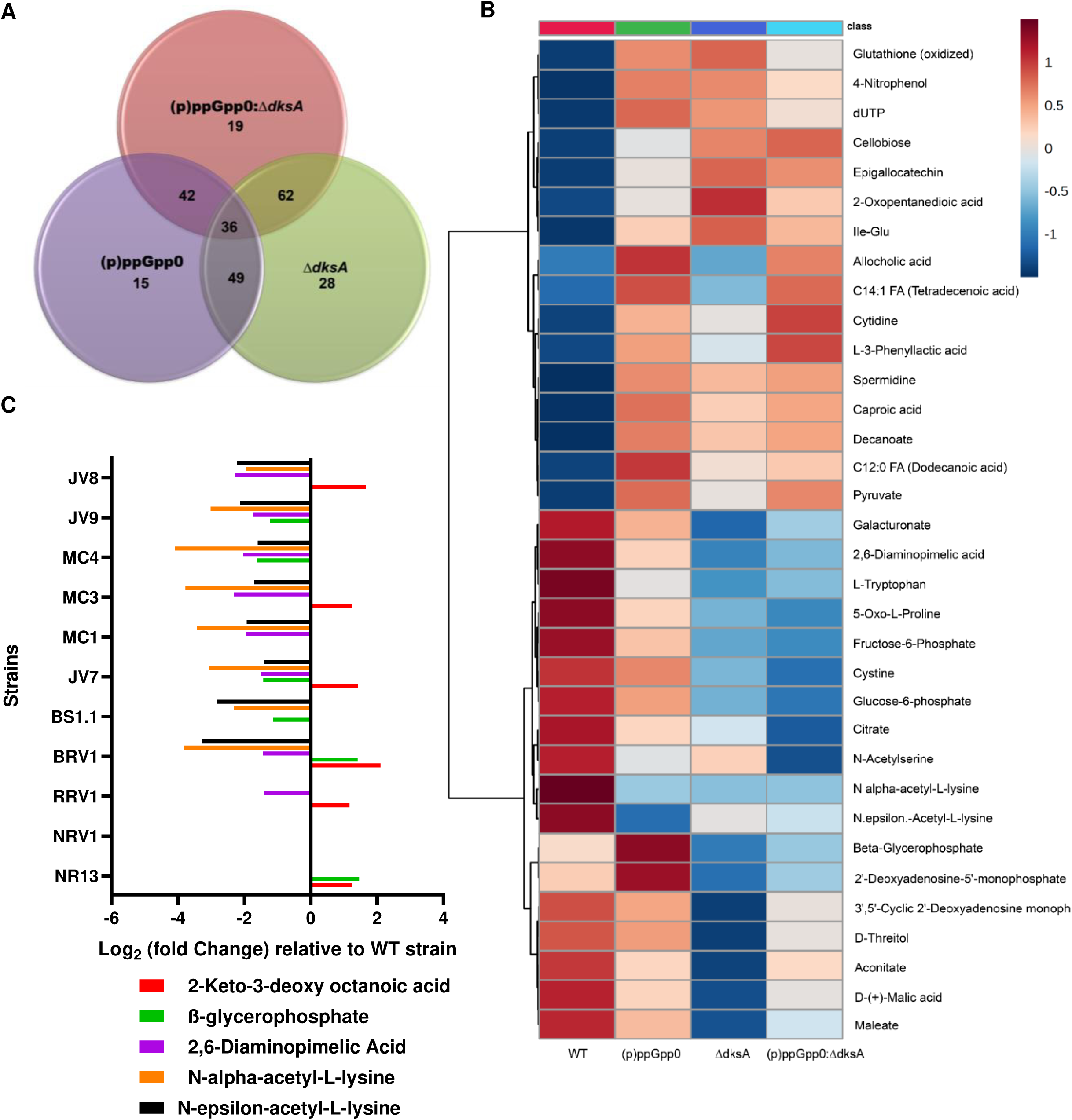
**A)** Venn diagram representing the numbers of altered metabolites in the BRV1, JV7 and JV9 mutant strains. 36 differential metabolites are overlapped among the three strains. **B)** Heatmap analysis of 36 overlapped metabolites among the BRV1, JV7 and JV9 strains. **C)** Log2 of the fold-change in the five important cell wall and membrane differentiated metabolites in the mutant strains compared with the WT (N16961) strain.

Subsequently, several metabolites were observed to be regulated by both (p)ppGpp alarmone and DksA transcription factor, either through collaborative or antagonistic mechanisms. The metabolites of BRV1, JV7 and JV9 strains are depicted in Venn diagram (Fig 4A). A total of 36 significantly altered metabolites were discovered to be shared by BRV1, JV7 and JV9 strains of *V. cholerae* and these metabolites were closely investigated. Heatmap analysis of 36 overlapped metabolites is shown in Fig 4B. Among the shared metabolites, few cell wall and membrane specific metabolites were observed to be differently regulated, among these include glycerol-2-phosphate (also known as β-glycerophosphate), involved in the synthesis of cell membrane. In contrast to the BRV1 strain, β-glycerophosphate was shown to be down regulated in JV7 and JV9 strains, illustrating antagonistic regulation. Whereas in BS1.1 strain β-glycerophosphate was observed to be downregulated, no change in the level of β-glycerophosphate in the JV8 strain. Apart from this 2,6-diaminopimelic acid (DAP), a major metabolite that is frequently found in the peptide links of NAM-NAG chains that constitute the cell wall of Gram-negative bacteria, was shown to be drastically downregulated in JV7, BRV1, and JV9 strains. Bacteria with a DAP shortage nevertheless grow normally but are not able to produce new peptidoglycan for their cell walls (28, 29). Also, N-α-acetyl-L-lysine and N-ε-acetyl-L-lysine, which is necessary for bacteria to produce lysine for protein synthesis, and cell wall biosynthesis, was similarly downregulated in all mutant strains. The formation and integrity of cell wall and membrane is compromised, suggesting a membrane re-composition linked to the basal levels of (p)ppGpp and presence and absence of DksA.

### Whole-cell proteome of (p)ppGpp^0^, Δ*dksA*, and (p)ppGpp^0^Δ*dksA* strains showed a reduced level of cell-wall biosynthetic enzymes

The regulatory roles of (p)ppGpp and DksA in influencing various cell wall associated biochemical pathways have been elucidated through metabolomic analyses. To enhance the mechanistic understanding of these regulatory processes, we explored the whole-cell proteome of mutant strains BRV1, JV7 and JV9. Utilizing the SWATH-MS platform, a total of 707, 731, and 684 proteins were quantified in BRV1, JV7 and JV9 strain, respectively, in comparison to the WT strain (Fig 5A). Several proteins involved in cell wall synthesis, organization, peptidoglycan biosynthesis, phospholipid transport, lipopolysaccharide transport and diaminopimelate biosynthetic processes, exhibited differential regulation in mutant strains (Fig 5B).

**Figure 5:**
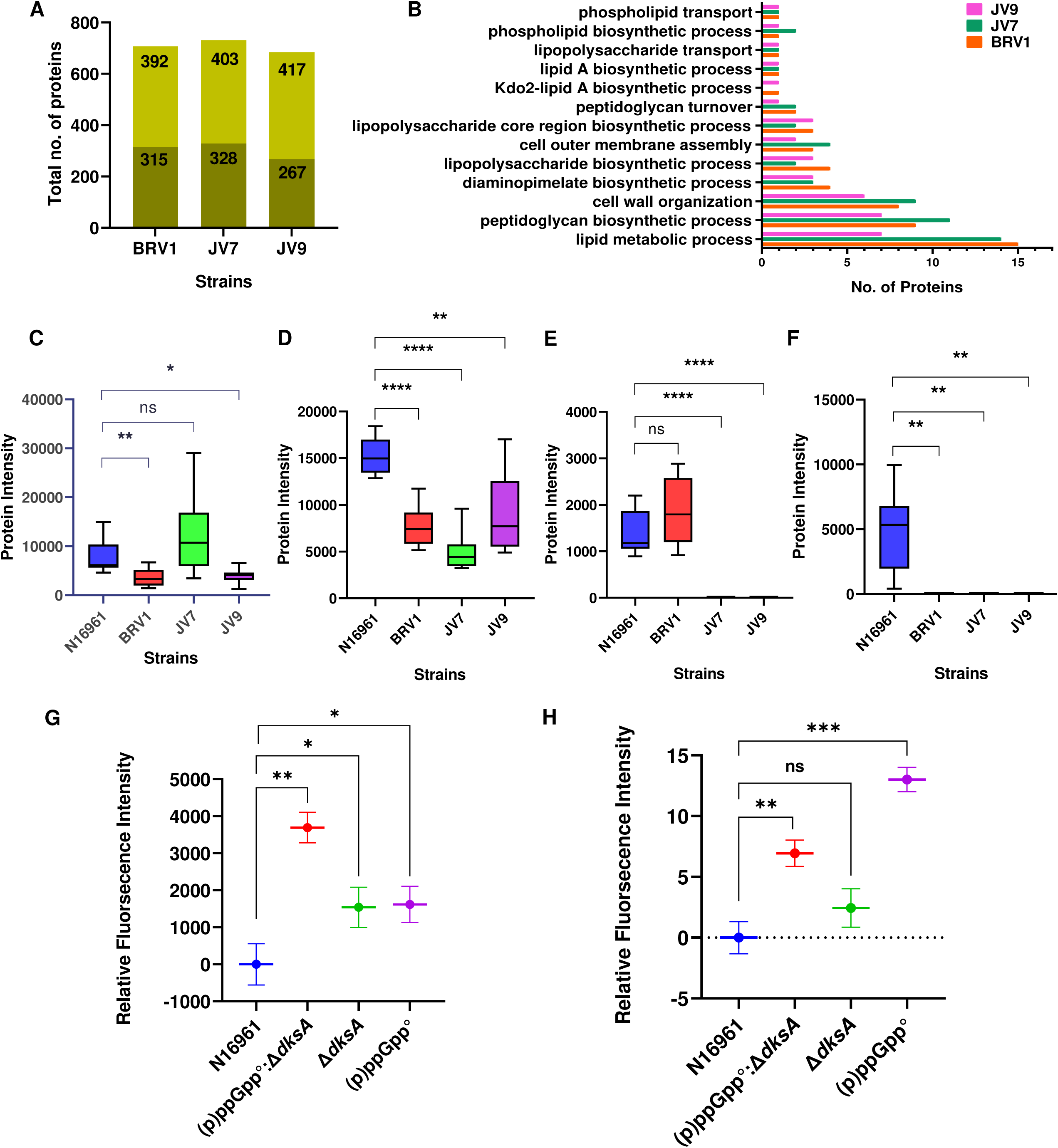
**A)** Total number of proteins identified by qualitative (lighter region) and quantitative analysis (darker region with fold-change ≥ 1.5 wrt WT) in BRV1, JV7 and JV9 strains. **B)** Comparison of cell wall and membrane associated pathways of differentially regulated proteins (mutant vs. WT) in BRV1, JV7 and JV9 strain. **C-F)** Box plot of protein intensities (mean area under the curve) for N16961, BRV1, JV7 and JV9 strains. Each plot represents a different protein. (**C**-PBP5, **D**-OmpU, **E**-LpoA, **F**-UDP-MurNAc-L-Ala-D-Glu:meso-diaminopimelate ligase.) [Welch’s t-test, (p < 0.05)] **G-H)** Relative fluorescence intensity for outer and inner membrane permeability assay detected by NPN and PI uptake, respectively in mutant strains.

The Penicillin-binding protein activator LpoA (PBP activator LpoA) was qualitatively identified in the WT and BRV1 strains but remained undetected in JV7, and JV9 strains. Furthermore, serine-type D-Ala-D-Ala carboxypeptidase (PBP5), a penicillin-sensitive, membrane-bound enzyme associated with resistance to β-lactam antibiotics, was observed to be down-regulated in BRV1(FC of 0.46) and JV9(FC of 0.512) strain. The enzymes involved in the DAP biosynthetic pathway exhibited distinct regulatory patterns in mutant strains. Notably, UDP-MurNAc-L-Ala-D-Glu:meso-diaminopimelate ligase, the enzyme responsible for integrating meso-diaminopimelic acid into the nucleotide precursor UDP-N-acetylmuramoy-L-alanyl-D-glutamate (UMAG), was qualitatively identified in the WT strain but remained undetected in BRV1, JV7, and JV9 strains. DAP decarboxylase, which catalyzes the final step of the DAP pathway, was also significantly downregulated in BRV1(FC of 0.604) and JV7(FC of 0.452) strains. This alignment with metabolomics data provides additional layers of validation, reinforcing the impact of (p)ppGpp and DksA on cell wall synthesis and dynamics by regulating the DAP pathway. In addition to this, outer membrane protein U (OmpU), was observed to be downregulated in BRV1(FC of 0.5), JV7 (FC of 0.321) and JV9 (FC of 0.585) strains (Fig 5C-F). Other than cell wall and membrane related proteins, we have also found numerous proteins showing differential expression in mutant strains in comparison to WT strain. These differentially expressed proteins predominantly belong to pathways involving metabolism, replication and translational regulation, and stress response. These findings strongly suggest the modulation of additional cellular pathways, which may be directly and/or indirectly linked with the increased susceptibility of antibiotics in (p)ppGpp and DksA mutant strains.

### Increased membrane permeability of (p)ppGpp^0^, and (p)ppGpp^0^Δ*dksA V. cholerae* strains

Metabolomic and proteomic analyses uncovered compelling evidence suggesting that (p)ppGpp and DksA affects the bacterial cell membrane synthesis which in turn impacts the structural and functional integrity of the cell membrane. Based on these findings, we have hypothesized that (p)ppGpp and DksA mutant strains may exhibit alterations in membrane permeability. To determine the effect of (p)ppGpp and DksA on inner membrane permeability (IMP) and outer membrane permeability (OMP), the assay using specific dyes was performed in BRV1, JV7 and JV9 strains. The OMP was evaluated by using NPN uptake assay. NPN is a neutral hydrophobic fluorescent probe which is usually excluded by the OM but when it partitions into the OM, it demonstrates increased fluorescence intensity. NPN fluorescence was significantly enhanced to the same level in BRV1, and in the JV9 strain however no significant difference was observed in the JV7 strain. The IMP was assessed by PI uptake assay as PI is a membrane-impermeant stain, only labels the bacteria having compromised IM. In contrast to OMP, the IMP was significantly enhanced in BRV1 and JV7 strains and a markedly enhanced effect was observed in JV9 (Fig 5G-H).

## DISCUSSION

Our previous analysis and other studies showed that the (p)ppGpp and DksA play crucial roles in stringent responses and help microbes to survive and overcome nutrient limitations and other environmental stress (8, 12). DksA and (p)ppGpp direct cellular resources from growth and multiplication processes to survival associated metabolic activities. However, intriguing disparities emerge at both the genotypic and phenotypic levels (17, 30). Stringent response modulators have been reported to modulate various cellular processes in Gram-negative bacteria such as fatty acid metabolism, amino acid metabolism, and flagellar synthesis, thereby determining virulence, pathogenicity, and bacterial survival under stressful conditions (31–34). The current study pursues to address the intricate interplay between (p)ppGpp and DksA in the modulation of antibiotic resistance in the absence of antibiotic resistance gene. The investigation spans phenotypic, metabolomic, and proteomic analyses in (p)ppGpp and DksA deficient strains providing a comprehensive understanding of the potential regulatory mechanisms underlying the increased antibacterial susceptibility in these mutant bacterial strains.

At phenotypic levels, we have observed that (p)ppGpp- and DksA-deficient strains of *V. cholerae* JV9, exhibit enhanced susceptibility to aminoglycoside, macrolide and β-lactam antibiotics. Notably, the most pronounced effects were observed for β-lactam antibiotics. The survival assay with β-lactam antibiotic ampicillin, imipenem, and aztreonam further underscored the susceptibility of (p)ppGpp- and DksA-deficient strains. In the absence of the β-lactam resistance genes and in the presence of sub-lethal concentrations of β-lactam antibiotics, survival for (p)ppGpp and DksA deficient strains were significantly diminished compared to WT strain and strain with higher levels of (p)ppGpp. Our findings aligned with earlier studies conducted in *E. coli* that have highlighted that (p)ppGpp-deficient (Δ*rel*AΔ*spo*T) mutant was more susceptible to cell-wall acting and other different classes of antimicrobial agents than the WT (19). The *ΔdksA* strain has also been demonstrated in *E. coli* to be more sensitive to antimicrobial drugs such as β-lactams, aminoglycosides, quinolones, and tetracyclines relative to the WT strain (20). This susceptibility pattern is also observed in *Acinetobacter baumannii,* where both (p)ppGpp-deficient and DksA deficient strains exhibited increased susceptibility to antimicrobial agents (18, 22). Despite the conventional application for targeting Gram-positive bacteria, vancomycin has also been included in our investigation as it has been extensively studied for its impact on (p)ppGpp-deficient strains in bacteria such as *Enterococcus faecalis, Staphylococcus aureus, Enterococcus faecium,* and *Bacillus subtilis* (35–37). We observed that survival is markedly reduced in (p)ppGpp and DksA deficient strains of *V. cholerae* in presence of sublethal concentration of vancomycin.

The mass spectrometry-based profiling method was conducted to study the level of metabolites in genetically engineered *V. cholerae* strains. In our current investigation, we observed that 50% of the differentially regulated metabolites were downregulated in both BRV1 and JV7 strains. Additionally, in the JV9 strain, 58% of the differently regulated metabolites exhibited downregulation. This highlights a predominant trend of metabolic alterations associated with (p)ppGpp and DksA in *V. cholerae*. Within the realm of membrane-specific metabolites, distinct patterns emerged. The UDP-GalNAc and KDO, which are precursors for Kdo2-lipid A, a principle and essential component of the OM of Gram-negative bacteria, are observed to be upregulated in BRV1 strain. Glycerol-2-phosphate displayed an antagonistic regulation, with elevated levels in the BRV1 strain and decreased levels in both the JV7 and BS1.1 strains. Furthermore, the absence of single or both SR modulators resulted in the downregulation of DAP levels. The previous transcriptional study conducted in *E. coli* has highlighted the substantial differences in the regulation of a significant number of genes involved in cell wall maintenance, plasma membrane function, and lipopolysaccharide (LPS) metabolism (15). Our current research aligns with prior findings, emphasizing that (p)ppGpp and DksA play a significant impact in shaping the dynamics of amino acid, nucleotide, and carbohydrate metabolism (6, 38).

To further confirm the findings of our phenotypic and metabolomics study, the total proteome of BRV1, JV7 and JV9 in comparison to WT strain was explored. The proteomic analysis revealed that the enzymes involved in purine salvage pathway (GuaB), translation (initiation factor 2 - IF2), and ribosome biogenesis (BipA, and RsgA) are significantly upregulated in BRV1 strain. Previous studies have reported these enzymes as a direct inhibitor of (p)ppGpp (39,40). Furthermore, differential regulation of various cell membrane and PG-associated proteins in the mutant strains was observed. The JV9 strain exhibits reduced expression of LpoA, PBP5, and OmpU, suggesting alterations in cell wall integrity and permeability of the outer membrane synthesis. LpoA has been reported to be essential for transpeptidase function of penicillin-binding protein 1A (PBP1a) and also to activate the upstream step of peptidoglycan polymerization (41). This indicates that the transpeptidase function of PBP1a and peptidoglycan polymerization might be indirectly linked to the expression of (p)ppGpp in combination with DksA. In line with the findings, the study in *E. coli* has highlighted that deletion of PBP5 makes the bacteria four- to eight-fold more susceptible to β-lactam antibiotics (42). Additionally, the recent study has highlighted the impact of (p)ppGpp on LPS biosynthesis by inhibiting LpxA, the first enzyme involved in LPS biosynthesis pathway (43). However, in our study no significant influence on LpxA was observed. The reliability of our metabolomics and proteomics findings is strengthened by the observed increased permeability of both IM and OM in the mutant strains.

In summary, our integrated genetic, proteomics, and metabolomics approach unveils a distinct regulatory network orchestrated by (p)ppGpp and DksA, elucidating their influence on reduced susceptibility to β-lactam antibiotics. The study highlighted the several causes of increased susceptibility to β-lactam in mutant strains, including modulation in cell membrane and PG synthesis activity, reduction in expression of PBPs and OMPs (Fig 6). Future work should focus on the development of inhibitors targeting SR modulators, with the aim of creating adjuvants that enhance the efficacy of β-lactam antibiotics.

**Figure 6:**
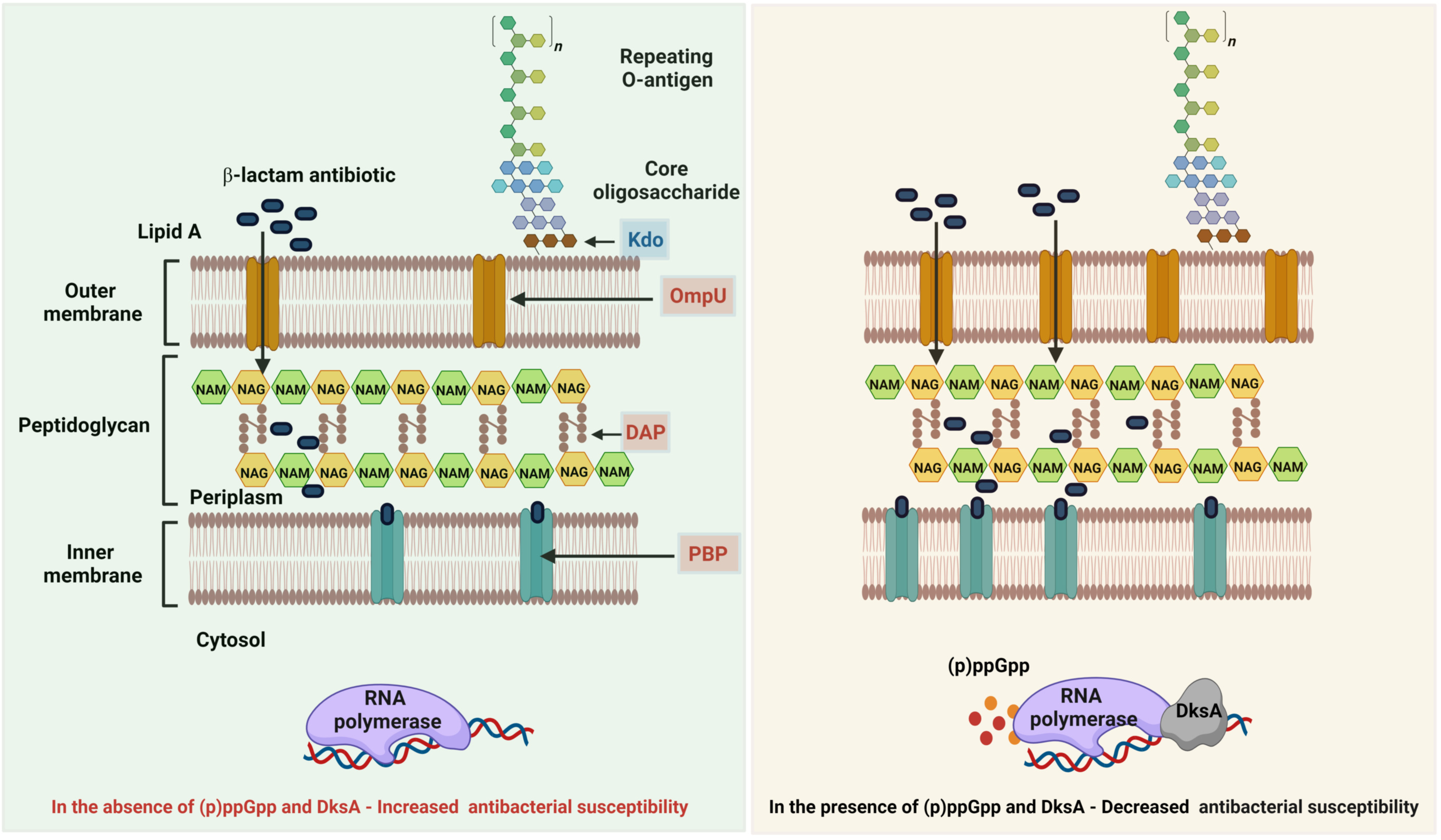
Schematic representation illustrating the heightened susceptibility to β-lactam antibiotics in (p)ppGpp and DksA mutant strain of *V. cholerae*. The increased susceptibility is attributed to distinct factors, including i) decreased expression of Penicillin-binding protein 5 (PBP5), ii) decreased expression of Outer membrane protein U (OmpU), iii) upregulation of 2- Keto-3-deoxy octanoic acid (Kdo), and iv) downregulation of 2,6-Diaminopimelic acid (DAP), indicating altered regulation of cell wall integrity and permeability of cell membrane in the mutant strains.

## MATERIALS AND METHODS

### Bacterial strains and growth conditions

The WT *V. cholerae* strain N16961 and previously constructed (p)ppGpp variant strains of *V. cholerae* and other strains constructed and used in this study are mentioned in **Table 1**. (p)ppGpp variant strains used were N16961-R13(N16961::Δ*relA*), N16961-RV1(N16961::Δ*relV*), BS1.1 (N16961::Δ*relA*, Δ*spoT*) and BRV1(N16961::Δ*relA*, Δ*spoT*, Δ*relV*) (11, 12). We have constructed DksA mutant strains of N16961 and of (p)ppGpp variant strains. All the plasmids used in this study are mentioned in Table S2. For liquid culture, the strains were grown in Luria Broth (LB) at 37 °C in a shaker with 180 rpm while LB agar plates were used for solid culture. The following antibiotic concentrations were used: streptomycin (100μg/mL), spectinomycin (50μg/mL), kanamycin (40μg/mL), zeocin (25μg/mL), ampicillin (100μg/mL) and chloramphenicol (30μg/mL for *E. coli* and 2μg/mL for *V. cholerae*). The bacteria were tested for sucrose sensitivity by plating them onto LA supplemented with 15% sucrose and incubating them at 24 °C. For long term storage at –80 °C, we used LB supplemented with 20% glycerol.

### Molecular biological methods

Unless otherwise specified, conventional molecular biology procedures were used for chromosomal and plasmid DNA isolations, electroelution, restriction enzyme digestion, ligation of DNA fragments, bacterial transformation, conjugation, and so on. All restriction enzymes and nucleic acid-modifying enzymes were sourced from New England BioLabs, Inc. and applied in accordance with manufacturer’s instructions. Transformants were chosen by plating transformed cells on LB agar plates with antibiotics.

### Development of recombinant vectors and mutant strains

Gene deletions and replacements were carried out utilizing the allelic exchange technique using suicide vector pDS132 derivatives. Using specific primer combinations, 500- to 700-base pair (bp) homologous areas upstream and downstream of the corresponding ORF were PCR-amplified (Table S3). The amplified components were purified even before restriction enzyme digestion, and ligated into a likewise digested suicide vector. The host bacterium *E. coli* FCV14 was utilized to select and replicate recombinant vectors. The recombinant vectors were transferred to *V. cholerae* via conjugation or electroporation. For conjugation, we used *E. coli* β-2163 as donor. The mutants were selected on respective antibiotics and further confirmed by PCR.

### Antibiotic susceptibility testing

Antibiotic susceptibility test by disc diffusion method was done to measure the zone of inhibition by different antibiotics in all (p)ppGpp and DksA mutant strains. For the disc diffusion method, all the strains were grown overnight aerobically at 37°C in MHB medium and the primary cultures were diluted 1:100 in fresh MHB medium and incubated aerobically at 37°C, when OD_600_ reached 0.5. The 1 ml of this culture was plated onto Mueller-Hinton agar (MHA, Difco, USA) plate (23” x 23” cm) using sterile cotton swabs and commercially available discs (Liofilchem) containing defined amounts of interested antibiotics were placed on it. Plates were incubated overnight at 37 °C in a static incubator and the zone of clearance was measured with the help of antibiotic zone scale.

### Survival assay by CFU count method

Survival assay of *V. cholerae* strains N16961, BS1.1, BRV1, JV7, JV8, JV9, MC1, MC3, and MC4 after exposure to sublethal concentration of antibiotics was measured by determining the number of CFU/mL. Overnight cultures were diluted 100-fold in 10 mL of fresh medium and incubated at 37°C with shaking at 180 RPM/min to reach OD_600_ = 0.5 (∼2 × 10^8^ CFU/mL). Aliquots of 5mL were then transferred into two different 50ml tubes. Sublethal concentration of antibiotics was added in one tube while another tube having no antibiotics was used as control. Tubes were incubated with shaking at 37°C for 4 hrs. For the determination of CFU, 1 mL aliquots were removed at the indicated time and cells were harvested by spinning down (8000 rpm for 2 min) followed by resuspension in fresh medium. Culture was serially diluted and plated on LB agar having appropriate sublethal concentration of antibiotics. Four antibiotics were selected primarily acting on cell wall synthesis for survival assay.

### Formula used

**Percentage of Survival = (Survival with antibiotic)/(Survival without antibiotic)×100**

Here, Survival with antibiotic = Total CFU at t=4 with antibiotic treatment – Total CFU at t=0

Survival without antibiotic = Total CFU at t=4 without antibiotic treatment – Total CFU at t=0

### Sample preparation for whole cell metabolome analysis

For the study of total cellular metabolites, the WT strain N16961 and different (p)ppGpp synthetic and *dksA* gene deleted strains of *V. cholerae*, were grown in LB at 37°C. An overnight culture of 1 ml was pelleted down by centrifugation ((10,000 rpm at 4°C for 10 min; Eppendorf centrifuge 5810R) and washed thrice with M9 minimal (M9M) medium. Culture was then diluted 100-fold in fresh casamino acid glucose-M9M medium and grown to late log phase (OD_600_ = 1.0). The cells were pelleted down again by centrifugation (10,000 rpm at 4°C for 10 min), washed with 0.9% normal saline and stored at -80°C. To extract the intracellular metabolites cold 100% methanol was added (Sigma Aldrich; Cat no. 34860) followed by vortexing and bath sonication for 10 min (Bransonic^®^ Ultrasonic M Cleaning Bath 1510). The cell debris was pelleted down by centrifugation (10,000 rpm at 4°C for 10 min) and supernatant was collected in two separate microcentrifuge tubes (120µl each tube), vacuum dried (Thermo Scientific™ Savant™ SPD1010) and stored at -80°C. For the analysis of metabolites, the dried supernatant was dissolved in 60µl of 15% methanol or 50% acetonitrile (Cat no. 271004) followed by vortexing for 5 min and centrifuged (10,000 rpm for 10 min). The supernatant was collected in a separate sample vial (Supelco™ Analytical). A small amount of each sample was also used to make a pool. Samples were submitted to run in six sets for the mass spectrometry.

### Measurement of metabolites

Data acquired using an Orbitrap Fusion mass spectrometer (Thermo Scientific) in conjunction with a heated electrospray ion source. With minor adjustments, data gathering procedures were carried out in accordance with published methods (44, 45). In summary, mass resolution was retained at 120,000 for MS1 mode and 30,000 for MS2 acquisition. The data acquisition mass range was 60-900Da. UPLC ultimate 3,000 was used to separate extracted metabolites. Data were collected using a reverse phase (RP) and HILIC column for positive and negative ionization modes. HSS T3 was used in the RP column, while XBridge BEH Amide was used in the HILIC column (Waters Corporation). Solvent A consisted of 20 mM ammonium acetate (pH-9.0) water for polar compound separation, while mobile phase B consisted of 100% acetonitrile. At a flow rate of 0.35 ml/min, the elution gradient commences at 85% B and proceeds to 10% B over 14 minutes. Solvent A for the RP was water, and Solvent B was methanol, with 0.1% formic acid in each. At a flow rate of 0.3 ml/min, the elution gradient proceeds from 1% B to 95% B in 10 minutes. The injection volume of the sample was 5µl. A pool quality control (QC) sample was taken after every five samples to evaluate signal variation and drift in mass inaccuracy.

### Data processing and analysis

The Progenesis QI software (Water Corporation) for metabolomics was used to process all LC/MS obtained data using the default settings. The untargeted workflow of Progenesis QI was used to achieve retention time alignment, deconvolution, feature identification, and elemental composition estimations. For database search, the Progenesis QI Metascope plug was used for the in-house library with right mass, retention period, and fragmentation pattern. For added identification certainty, we employed an online spectrum library. The retention time match cut- off in Progenesis QI was 0.5 min, and spectral similarity was more than 30% fragmentation match. Peaks in pool QC samples with a coefficient of variation (CV) less than 30% were saved for further analysis. Furthermore, each found characteristic was thoroughly confirmed before being used to choose relevant peaks.

For the univariate and multivariate analysis performed with Metaboanalyst 5.0, the resulting data matrices were sum normalized, log transformed and Pareto scaled. Principal Component Analysis (PCA) was done to understand the clustering pattern. Metabolites with fold change threshold of 2 and above and metabolites passing FDR-adjusted-p-value were considered for the study. Metaboanalyst 5.0 enrichment analysis was used to evaluate the top chemical classes in mutant strains. Heatmap analyses were performed to determine features of statistical significance among mutant groups. Information from EcoCyc for metabolic pathway analysis was used.

### Whole cell proteome analysis

For the study of total cell proteins, the WT strain N16961 and BRV1, JV7 and JV9 mutants of *V. cholerae* were selected for proteome analysis. The organisms were grown in LB medium at 37°C with shaking at 180 rpm overnight. Overnight culture of 1 ml was pelleted down and washed thrice with M9M medium. Culture was then diluted 100-fold in fresh casamino acid glucose- M9M medium and grown to late log phase (OD_600_ = 1.0). The cells were pelleted down by centrifugation, washed with chilled PBS and whole-cell proteins were extracted using lysis buffer (8M urea, 2% sodium deoxycholate in 50mM ammonium bicarbonate, pH 7.2). Protein samples were digested into peptides using trypsin before reduction and alkylation with dithiothreitol and iodoacetamide, respectively. The resulting peptide mixtures were desalted and analyzed by Eksigent microLC, connected with a TripleTOF 5600 mass spectrometer. 10 µg desalted peptides were loaded on a reverse phase analytical column (3C18-CL, 300µm x 15mm, 3µm, 120Å) using mobile phases as follows: water/acetonitrile/formic acid (A, 98/2/0.2%; B, 2/98/0.2%). Separation was established by the following gradient condition: initial 5% B for 5 min, followed by a linear gradient from 3% B to 25% B in 68 min, followed by another linear gradient to 35% B in 73 min. Following the peptide elution window, the gradient was increased to 80% B in 2 min and held for 3 min. Initial chromatographic conditions were restored in 1 min and maintained for 8 min. DDA spectra were acquired using an ESI ion source with 100 – 1500 m/z mass range and cycle time 2.3s. A set of 83 overlapping SWATH windows were also used to acquire the data in DIA mode with a total duty cycle of 4.02s. Identification of proteins was carried out via search against Uniprot protein databases of *V. cholerae* with ProteinPilot to obtain a spectral library. The proteins and associated peptides were filtered by PeakView and MarkerView for relative quantitative and qualitative analysis. All the proteins with fold change threshold of 1.5 and above and p-value < 0.05 were considered for the study.

### Outer membrane permeability analysis

A single colony of WT and mutant strains of *V. cholerae* was inoculated into 5 ml LB broth in a glass culture tube and incubated overnight in a shaking incubator at 180 rpm at 37 °C. 1 ml of overnight culture was pelleted down and washed thrice with M9M medium. Culture was then diluted 100-fold in fresh casamino glucose-M9M medium and grown to late-log phage (OD_600_ = 1.0). The cells were pelleted down at 4°C and after washing twice with chilled PBS, resuspended in the same volume of PBS. 200µl was added to each 96-well plate. For outer membrane permeability, 1-N-phenylnaphthylamine (NPN) was used at a final concentration of 15µM. The reading was taken immediately at excitation wavelength of 350 nm, and emission wavelength of 420 nm. The experiment was done in triplicates.

### Inner membrane permeability assay

The bacterial cells were prepared as mentioned above in section 2.7. For inner membrane permeability, propidium iodide was used at a final concentration of 2.5µg/ml. The reading was taken at excitation wavelength of 535 nm, and emission wavelength of 617 nm. The experiment was done in triplicates.

---

**Supplementary Figure 1:** The graph represents the relative zone of Inhibition of *V. cholerae* WT, NRV1, NR-13, RRV1, MC4, MC1 and MC3 strains with different antibiotics.

**Supplementary Figure-2:** Volcano plot of **A)** BRV1, and **B)** JV7 strains of the 291 identified metabolites by LC–MS. The volcano plot shows the fold-change (x-axis) versus the significance (y-axis) of the 291 metabolites. The vertical and horizontal dotted lines show the cut-off of fold-change = ± 2, and of p-value = 0.05, respectively.

## Contributors

BD conceived the idea and designed the experiments. MC, JV, SK, TM, TS, PB, YK, conducted the experiments. RKB contributed reagents. MC, JV, BD, performed data analysis. MC and BD wrote the manuscript. RKB, and BD edited the manuscript. All authors have read and approved the manuscript.

## Acknowledgement

We acknowledge the generous support of Prof. G Karthikeyan and Prof. Pramod K Garg to complete the study. We thankfully acknowledge the Advanced Nucleotide Sequencing Facility at THSTI for gene and genome sequencing. The work was funded by the Translational Research Program (TRP) (No. BT/PR30159/MED/15/188/2018) of Department of Biotechnology (DBT), Govt. of India. RKB received support from the Emeritus Scheme Grant [21(1100)/20/EMR-II], Council of Scientific and Industrial Research (CSIR), Government of India. The funding agency had no role in study design, sample collection, analysis and interpretation of data and writing the manuscript. MC received research fellowship from the CSIR, Govt. of India (CSIR-File No:09/1049(0039)/2020-EMR-I). SK received research fellowship from The University Grants Commission (UGC), Govt. of India.

## References

1. Dalebroux ZD, Swanson MS. 2012. ppGpp: magic beyond RNA polymerase. Nat Rev Microbiol 10:203–212.

2. Hauryliuk V, Atkinson GC, Murakami KS, Tenson T, Gerdes K. 2015. Recent functional insights into the role of (p)ppGpp in bacterial physiology. Nat Rev Microbiol 13:298– 309.

3. Iyer S, Le D, Park BR, Kim M. 2018. Distinct mechanisms coordinate transcription and translation under carbon and nitrogen starvation in Escherichia coli. Nat Microbiol 3:741–748.

4. Srivatsan A, Wang JD. 2008. Control of bacterial transcription, translation and replication by (p)ppGpp. Curr Opin Microbiol 11:100–105.

5. Potrykus K, Cashel M. 2008. (p)ppGpp: still magical? Annu Rev Microbiol 62:35–51.

6. Irving SE, Choudhury NR, Corrigan RM. 2021. The stringent response and physiological roles of (pp)pGpp in bacteria. Nat Rev Microbiol 19:256–271.

7. Gourse RL, Chen AY, Gopalkrishnan S, Sanchez-Vazquez P, Myers A, Ross W. 2018. Transcriptional Responses to ppGpp and DksA. Annu Rev Microbiol 72:163–184.

8. Pal RR, Bag S, Dasgupta S, Das B, Bhadra RK. 2012. Functional characterization of the stringent response regulatory gene dksA of Vibrio cholerae and its role in modulation of virulence phenotypes. J Bacteriol 194:5638–5648.

9. Kang PJ, Craig EA. 1990. Identification and characterization of a new Escherichia coli gene that is a dosage-dependent suppressor of a dnaK deletion mutation. J Bacteriol 172:2055–2064.

10. Paul BJ, Barker MM, Ross W, Schneider DA, Webb C, Foster JW, Gourse RL. 2004. DksA: a critical component of the transcription initiation machinery that potentiates the regulation of rRNA promoters by ppGpp and the initiating NTP. Cell 118:311–322.

11. Burgos HL, O’Connor K, Sanchez-Vazquez P, Gourse RL. 2017. Roles of Transcriptional and Translational Control Mechanisms in Regulation of Ribosomal Protein Synthesis in Escherichia coli. J Bacteriol 199.

12. Das B, Pal RR, Bag S, Bhadra RK. 2009. Stringent response in Vibrio cholerae: genetic analysis of spoT gene function and identification of a novel (p)ppGpp synthetase gene. Mol Microbiol 72:380–398.

13. Das B, Bhadra RK. 2007. Molecular characterization of Vibrio cholerae ΔrelA ΔspoT double mutants. Arch Microbiol 189:227–238.

14. Dasgupta S, Das B, Basu P, Bhadra RK. 2016. Molecular Basis of the Stringent Response in Vibrio Cholerae, p. 507–516. In Stress and Environmental Regulation of Gene Expression and Adaptation in Bacteria. John Wiley & Sons, Ltd.

15. Sanchez-Vazquez P, Dewey CN, Kitten N, Ross W, Gourse RL. 2019. Genome-wide effects on transcription from ppGpp binding to its two sites on RNA polymerase. Proc Natl Acad Sci U S A 116:8310–8319.

16. Chen J, Gopalkrishnan S, Chiu C, Chen AY, Campbell EA, Gourse RL, Ross W, Darst SA. 2019. E. coli TraR allosterically regulates transcription initiation by altering RNA polymerase conformation and dynamics. bioRxiv.

17. Huang C, Meng J, Li W, Chen J. 2022. Similar and Divergent Roles of Stringent Regulator (p)ppGpp and DksA on Pleiotropic Phenotype of Yersinia enterocolitica. Microbiol Spectr 10:e0205522.

18. Kim N, Son J-H, Kim K, Kim H-J, Shin M, Lee J-C. 2021. DksA Modulates Antimicrobial Susceptibility of. Antibiotics (Basel) 10.

19. Greenway DL, England RR. 1999. The intrinsic resistance of Escherichia coli to various antimicrobial agents requires ppGpp and sigma s. Lett Appl Microbiol 29:323–326.

20. Wang J, Cao L, Yang X, Wu Q, Lu L, Wang Z. 2018. Transcriptional analysis reveals the critical role of RNA polymerase-binding transcription factor, DksA, in regulating multi- drug resistance of Escherichia coli. Int J Antimicrob Agents 52:63–69.

21. Viducic D, Ono T, Murakami K, Susilowati H, Kayama S, Hirota K, Miyake Y. 2006. Functional analysis of spoT, relA and dksA genes on quinolone tolerance in Pseudomonas aeruginosa under nongrowing condition. Microbiol Immunol 50:349–357.

22. Jung H-W, Kim K, Islam MM, Lee JC, Shin M. 2020. Role of ppGpp-regulated efflux genes in Acinetobacter baumannii. J Antimicrob Chemother 75:1130–1134.

23. Anderson MS, Bulawa CE, Raetz CR. 1985. The biosynthesis of gram-negative endotoxin. Formation of lipid A precursors from UDP-GlcNAc in extracts of Escherichia coli. J Biol Chem 260:15536–15541.

24. Bertani B, Ruiz N. 2018. Function and Biogenesis of Lipopolysaccharides. EcoSal Plus 8.

25. Belunis CJ, Raetz CR. 1992. Biosynthesis of endotoxins. Purification and catalytic properties of 3-deoxy-D-manno-octulosonic acid transferase from Escherichia coli. J Biol Chem 267:9988–9997.

26. Brozek KA, Hosaka K, Robertson AD, Raetz CR. 1989. Biosynthesis of lipopolysaccharide in Escherichia coli. Cytoplasmic enzymes that attach 3-deoxy-D-manno-octulosonic acid to lipid A. J Biol Chem 264:6956–6966.

27. Wachsmuth K. 1994. Vibrio Cholerae and Cholera: Molecular to Global Perspectives.

28. Pasternak CA. 1981. Microbial cell walls and membranes: By H. J. Rogers, H. R. Perkins AND J. B. Ward. 1980. Chapman and Hall, Ltd, London. Pp. viii and 564. 30.00. Journal of Medical Microbiology 10.1099/00222615-14-4-501.

29. Dörr T, Cava F, Lam H, Davis BM, Waldor MK. 2013. Substrate specificity of an elongation-specific peptidoglycan endopeptidase and its implications for cell wall architecture and growth of Vibrio cholerae. Mol Microbiol 89:949–962.

30. Łyżeń R, Maitra A, Milewska K, Kochanowska-Łyżeń M, Hernandez VJ, Szalewska-Pałasz A. 2016. The dual role of DksA protein in the regulation of Escherichia coli pArgX promoter. Nucleic Acids Res 44:10316–10325.

31. Dasgupta S, Das S, Biswas A, Bhadra RK, Das S. 2019. Small alarmones (p)ppGpp regulate virulence associated traits and pathogenesis of Salmonella enterica serovar Typhi. Cell Microbiol 21:e13034.

32. Aberg A, Fernández-Vázquez J, Cabrer-Panes JD, Sánchez A, Balsalobre C. 2009. Similar and divergent effects of ppGpp and DksA deficiencies on transcription in Escherichia coli. J Bacteriol 191:3226–3236.

33. Zhang Y, Teper D, Xu J, Wang N. 2019. Stringent response regulators (p)ppGpp and DksA positively regulate virulence and host adaptation of Xanthomonas citri. Mol Plant Pathol 20:1550–1565.

34. Huang C, Li W, Chen J. 2023. Transcriptomic Analysis Reveals Key Roles of (p)ppGpp and DksA in Regulating Metabolism and Chemotaxis in. Int J Mol Sci 24.

35. Singh M, Matsuo M, Sasaki T, Morimoto Y, Hishinuma T, Hiramatsu K. 2017. In Vitro Tolerance of Drug-Naive Staphylococcus aureus Strain FDA209P to Vancomycin. Antimicrob Agents Chemother 61.

36. Corrigan RM, Bellows LE, Wood A, Gründling A. 2016. ppGpp negatively impacts ribosome assembly affecting growth and antimicrobial tolerance in Gram-positive bacteria. Proc Natl Acad Sci U S A 113:E1710–9.

37. Abranches J, Martinez AR, Kajfasz JK, Chávez V, Garsin DA, Lemos JA. 2009. The molecular alarmone (p)ppGpp mediates stress responses, vancomycin tolerance, and virulence in Enterococcus faecalis. J Bacteriol 191:2248–2256.

38. Durfee T, Hansen A-M, Zhi H, Blattner FR, Jin DJ. 2008. Transcription profiling of the stringent response in Escherichia coli. J Bacteriol 190:1084–1096.

39. Kriel A, Bittner AN, Kim SH, Liu K, Tehranchi AK, Zou WY, Rendon S, Chen R, Tu BP, Wang JD. 2012. Direct regulation of GTP homeostasis by (p)ppGpp: a critical component of viability and stress resistance. Mol Cell 48:231–241.

40. Zhang Y, Zborníková E, Rejman D, Gerdes K. 2018. Novel (p)ppGpp Binding and Metabolizing Proteins of. MBio 9.

41. Hunashal Y, Kumar GS, Choy MS, D’Andréa ÉD, Da Silva Santiago A, Schoenle MV, Desbonnet C, Arthur M, Rice LB, Page R, Peti W. 2023. Molecular basis of β-lactam antibiotic resistance of ESKAPE bacterium E. faecium Penicillin Binding Protein PBP5. Nat Commun 14:4268.

42. Sarkar SK, Chowdhury C, Ghosh AS. 2010. Deletion of penicillin-binding protein 5 (PBP5) sensitises Escherichia coli cells to beta-lactam agents. Int J Antimicrob Agents 35:244–249.

43. Brückner S, Müller F, Schadowski L, Kalle T, Weber S, Marino EC, Kutscher B, Möller A-M, Adler S, Begerow D, Steinchen W, Bange G, Narberhaus F. 2023. (p)ppGpp and moonlighting RNases influence the first step of lipopolysaccharide biosynthesis in. Microlife 4:uqad031.

44. Naz S, Gallart-Ayala H, Reinke SN, Mathon C, Blankley R, Chaleckis R, Wheelock CE. 2017. Development of a Liquid Chromatography-High Resolution Mass Spectrometry Metabolomics Method with High Specificity for Metabolite Identification Using All Ion Fragmentation Acquisition. Anal Chem 89:7933–7942.

45. Kumar A, Kumar Y, Sevak JK, Kumar S, Kumar N, Gopinath SD. 2020. Metabolomic analysis of primary human skeletal muscle cells during myogenic progression. Sci Rep 10:11824.

